# Temporal Small RNA Expression Profiling Under Drought Reveals a Potential Regulatory Role of snoRNAs in Drought Responses of Maize

**DOI:** 10.1101/376004

**Authors:** Jun Zheng, Erliang Zeng, Yicong Du, Cheng He, Ying Hu, Zhenzhen Jiao, Kai Wang, Wenxue Li, Maria Ludens, Junjie Fu, Haiyan Wang, Frank F. White, Guoying Wang, Sanzhen Liu

## Abstract

Small RNAs (sRNAs) are short noncoding RNAs that play roles in many biological processes, including drought responses in plants. However, how the expression of sRNAs dynamically changes with the gradual imposition of drought stress in plants is largely unknown. We generated time-series sRNA sequence data from maize seedlings under drought stress and under well-watered conditions at the same time points. Analyses of length, functional annotation, and abundance of 736,372 non-redundant sRNAs from both drought and well-watered data, as well as genome copy number and chromatin modifications at the corresponding genomic regions, revealed distinct patterns of abundance, genome organization, and chromatin modifications for different sRNA classes of sRNAs. The analysis identified 6,646 sRNAs whose regulation was altered in response to drought stress. Among drought-responsive sRNAs, 1,325 showed transient down-regulation by the seventh day, coinciding with visible symptoms of drought stress. The profiles revealed drought-responsive microRNAs, as well as other sRNAs that originated from ribosomal RNAs (rRNAs), splicing small nuclear RNAs, and small nucleolar RNAs (snoRNA). Expression profiles of their sRNA derivers indicated that snoRNAs might play a regulatory role through regulating stability of rRNAs and splicing small nuclear RNAs under drought condition.

## Introduction

Physiological responses to drought in plants are complex and regulated through an interplay of a network of genetic components and chromatin structure. One component is comprised of drought-responsive small RNAs (sRNAs) (Khraiwesh *et al*. 2012). sRNAs are short noncoding RNAs, predominately 20 to 24 nt in length, which function as sequence-specific regulators in a wide variety of biological processes, including DNA methylation, RNA degradation, translation regulation, and histone modification (Khraiwesh *et al*. 2012; Axtell 2013a). Plant sRNAs are typically categorized into two major groups, which are distinguished by the structure of the sRNA precursors. The first group consists of microRNAs (miRNAs), which are predominately 21 nt in length and processed from single-stranded precursor RNA, or pri-sRNA, are transcribed by RNA polymerase II (PolII) and contain a hairpin structure. The second group is comprised of small interfering RNAs (siRNAs) are derived from DICER/DICER-like processing of double-stranded RNAs (dsRNAs).

MiRNAs function in drought stress responses (Covarrubias and Reyes 2010; Shuai *et al*. 2013) and are, conceptually, categorized into three functional categories - homeostasis, detoxification, and growth regulation (Zhu 2002) and function largely through the destabilization of various transcription factors (Rhoades *et al*. 2002; Ding *et al*. 2013; Ferdous *et al*. 2015; Zhang 2015). The function of miRNAs in the regulation of transcription factors places miRNAs at the hubs of gene regulatory networks for drought responses. whereas miRNAs primarily act in the posttranscriptional regulation of gene expression, siRNAs regulate gene transcription through both guiding DNA methylation by the pathway of RNA-directed DNA methylation (RdDM) and posttranscriptional destabilization of transcripts in a sequence-specific manner (Onodera *et al*. 2005; Wierzbicki *et al*. 2008). Small interfering RNAs can be further sub-grouped into heterochromatic siRNAs, secondary siRNAs, and natural antisense transcripts siRNAs (NAT-siRNA). Heterochromatic siRNAs, typically, are 23-24 nt in length and require RNA-dependent RNA polymerase (RDR) and RNA polymerase IV (PolIV) for biogenesis. Heterochromatic siRNAs were documented to be derived from transposable/repetitive elements located at heterochromatic regions of nuclear DNA (Meyers *et al*. 2008; Nobuta *et al*. 2008). Secondary siRNAs include trans-acting siRNAs (ta-siRNA), which are formed through cleavage of capped and polyadenylated siRNA transcripts by specific miRNAs, followed by conversion into dsRNAs by RDR (Vazquez *et al*. 2010). NAT-siRNAs are derived from dsRNAs formed by annealing of natural sense and antisense transcripts from the same or separate nearly identical genomic regions (Vazquez *et al*. 2010).

Small RNAs can also originate from ribosomal RNAs (rRNAs), transfer RNAs (tRNAs), small nucleolar RNAs (snoRNAs), and small nuclear RNAs associated with mRNA splicing (splicing snRNAs) and are respectively referred to as rsRNAs, tsRNAs, sno-sRNAs, and splicing sn-sRNAs hereafter (Vazquez *et al*. 2010). The rsRNAs, tsRNAs, sno-sRNAs, and splicing sn-sRNAs play regulatory roles in cellular processes (Morris and Mattick 2014). In barley, tsRNAs and sno-sRNAs tended to be up-regulated and down-regulated, respectively, under drought conditions (Hackenberg *et al*. 2015). In maize, miRNA biosynthesis and regulation under drought stress has been explored (LI *et al*. 2013; Liu *et al*. 2014; Wang *et al*. 2014). However, the regulatory functions of sRNAs other than miRNAs are largely unknown.

To understand sRNA function and regulation in the drought response of maize, we sequenced sRNAs from maize seedlings over a period of 3 to 11 days after witholding water along with sRNAs from well-watered plants or drought treated plants that recovered after watering. The sRNAs were categorized with respect to length and functional classification, and the genomic organization of sRNAs was analyzed. An attempt was made to classify drought-responsive sRNAs using cluster and network analyses on the time-series expression patterns, providing clues of destabilization of ribosome RNA and splicing small nuclear RNAs under drought condition.

## Materials and Methods

### Plant materials and drought treatments

Seeds of the maize (*Zea mays*) inbred line B73 were surface-sterilized and germinated on the wet rolled brown paper towel at 28°C for 48h, and eighteen germinated seeds were selected and transplanted in a plastic pot (17×12×10cm) filled with nutrient soil (1:1 peat moss and vermiculite). Three-day seedlings after germination were subjected to drought stress up to 10 days by withholding water (10 DAW), and the control plants were well watered. The plants were grown on the controlled conditions (27 °C day/23 °C night, 16 h photoperiod, from 6 am to 10 pm, 300μmol m^−2^ s^−1^ photons, 30-50% relative humidity). Treatment (drought stress) and the control pots were randomly laid in growth chamber. Eighteen seedlings were planted in a pot. For every harvest and sample time, five pots were used for a drought treatment and other 5 pots were used as a control. At 10 DAW, drought treated seedling plants were divided into two groups: a group of seedlings kept under drought stress without watering and the other group of seedlings that were re-watered. In summary, 36 samples of soils and leaf tissues were collected: i) 32 samples resulting from 2 treatments (drought stress (DS) and well watered (WW)) x 8 days (from day 3 to day 10) x 2 biological replicates; ii) 4 samples resulting from 2 treatments (DS and WW at day 11th of plants previously subjected to 10 days of DS) x 2 biological replicates.

### Measurement of soil SWC, leaf RWC, and leaf REC

Soil samples and leaf tissues for measuring SWC (soil water content), RWC (relative water content), and REC (Relative electrical conductivity) were daily collected at around 9:30 am. Five independent replicates were performed for the SWC measurement, and five biological replicates were performed for RWC and REC measurements. SWC, RWC, and REC were carried out according to the previously described method (Zheng *et al*. 2010). Briefly, the soil SWC was the percentage of the weight loss of soils after drying. The RWC of the fresh leaves was calculated using the formula of (FW-DW)/(TW-DW) x100%, where FW is the weight of fresh leaves, TW is the leaf weight after saturated in water for 8 h, and DW is the leaf dry weight. REC was calculated using Ec1/Ec2 x 100%, where Ec1 is electrical conductivity of fresh leaves after saturated in water for 3 h and Ec2 is electrical conductivity of the same leaf samples after boiled in a water bath.

### sRNA sequencing experiment

The above ground tissues of five seedlings of each treatment at each day were collected at approximately 10 am each day and immediately frozen in liquid nitrogen. Total RNA was isolated from harvested samples using TRIzol reagent (Invitrogen). A standard Illumina small RNA library preparation kit was used to prepare small RNA sequencing libraries from total RNAs. Briefly, a total of 2 μg sRNAs in the size range of 15 to 30 nucleotides were purified and ligated to 3’ adaptor, and isolated by 15% denaturing polyacrylamide gel electrophoresis gels to eliminate un-ligated 3’ adaptors. The products were ligated to 5’ adaptor and then were used to conduct reverse transcription PCR. The final PCR product was isolated by 3.5% agarose gel electrophoresis and served as a small RNA library for the sequencing. The libraries were quantified and sequenced at HiSeq2000 analyzer to produce single-end 50 bp reads. Two biological replicates were employed in the sRNA sequencing experiment.

### sRNA data process

Trimmomatic (version 0.32) was used to trim the adaptor sequence of sRNA reads. The parameters used for the trimming is: “ILLUMINACLIP:adaptor_seq:2:30:7: LEADING:3 TRAILING:3 SLIDINGWINDOW:4:13 MINLEN:16”. The adaptor sequence (adaptor_seq) includes a sequence of “CTGTAGGCACCATCAATCAGATCGGAAGAGCACACGTCTGAACTCCAGTCAC”. These parameters were used to perform both adaptor and quality trimming. Although quality trimming could shorten actual sRNAs, the percentage of reads subjected to quality trimming is only ~0.3%. Therefore, quality trimming was applied to remove the low quality of nucleotides at the marginal compromise of changing sRNA lengths. At least 16 nt in size was required for clean reads.

A non-redundant sRNA (NR-sRNA) set was obtained by pooling sRNAs from all the samples and remove the redundancy. To remove most sRNA sequences that carry sequence errors, only sRNAs that were shown in at least two different samples and at least twice in each sample were included in the unique sRNA set. After determining read counts of each sRNA from all 36 samples, a further reduction was performed to only keep sRNAs with at least 72 reads summed from all the samples, equivalent to 0.08 reads per million of total reads, resulting in a NR-sRNA set.

### Functional annotation of sRNAs

The small RNA annotation database was downloaded from Rfam 11.0 (Burge *et al*. 2013). sRNAs generated from this experiment were aligned to Rfam 11.0 database using Blastn (BLAST 2.2.29+) with the following parameters (-evalue 1e-1 –word_size 10 – perc_identity 0.89 –strand plus –best_hit_overhang 0.2 –best_hit_score_edge 0.1 –outfmt 6 –max_target_seqs 10). The sRNAs was functionally annotated only if they were unambiguously hit an Rfam family.

### Alignment to the reference genome to determine copy number of sRNA regions

Each sRNA was aligned to the B73 reference genome (RefGen2 and 4) using bwa (version 0.7.5a-r405) (Li and Durbin 2010). The command parameters were “bwa aln –l 18 –k 0 –t 48 –R 22500” followed by “bwa samse –n 22500”. The alignments were then parsed with the stringent criteria: perfect match with at least 18 bp matching length. These alignment and parsing criteria allow the maximal 22,500 perfect hits.

### K-mer analysis using sequencing data to determine copy number of sRNA genomic regions

B73 whole genome shotgun Illumina sequencing data were downloaded from Genbank (SRR444422). Trimmomatic (version 0.32) was used for the adaptor and quality trimming with the same parameters to those used in the sRNA data trimming. The adaptor sequences used for the adaptor trimming are (TACACTCTTTCCCTACACGACGCTCTTCCGATCT and GTGACTGGAGTTCAGACGTGTGCTCTTCCGATCT). The clean data were subjected to error correction using the error correction module (ErrorCorrectReads.pl) in ALLPATHS-LG (Butler *et al*. 2008) with the parameters of “PHRED_ENCODING=33 PLOIDY=1”. We then used the corrected sequencing data to perform k-mer counting using the count function in JELLYFISH (Marcais and Kingsford 2011) with the parameters of “-m k-mer –L 2 –s 100M –C”, where the k-mer was from 18 to 30 nt. Once the read depth of each k-mer from 18 to 30 nt was counted, the read depth of a corresponding sRNA can be determined. The highest density of k-mer counts was located at 26.96 for a set of known single copy k-mers determined by reference genome alignments, indicating approximately 26.96x sequencing depth was obtained. This number was used as the base of read depths of a single copy to adjust counts of each k-mer to roughly represent its genome copy number.

### Determination of mean levels of various histone modifications

ChIP-Seq data of H3K27me3, H3K36me3, H3K4me3, and H3K9ac (Wang *et al*. 2009), and H3K9me2 (West *et al*. 2014) of 14-day B73 seedlings were downloaded from Genbank. To match sequencing data to sRNA sequences, ChIP-Seq data were subjected to k-mer counts at different k-mer lengths from 18 to 26 nt using JELLYFISH (Marcais and Kingsford 2011). Through k-mer counts, read counts from ChIP-Seq data of each sRNA sequence was determined. Using sequencing read counts of whole genome sequencing (WGS) data of sRNA sequences as the control, the histone modification signal of each sRNA, represented by ChIP read count divided by WGS read count of an sRNA, was calculated. Due to the lack of biological replication and the limited sequencing depth of ChIP-Seq data, we did not attempt to assess the histone modification level of each sRNA. Instead, mean of histone modification levels of all sRNAs in a certain functional group (e.g., miRNA, rsRNA) were determined and used as the modification level of that sRNA group for the comparison between functional groups. Comparisons were only performed within the same length of sRNAs.

To enable the comparison of histone modification levels among different lengths of sRNAs, all lengths of sRNAs from 18 to 26 nt were converted to 18 nucleotide fragments and the average signal of five epimarks were determined separately. As a control, 18 nt of different genic regions, promoters, first exons, internal exons, introns, and last exons, were sampled and the average histone modification levels of five epimarks of each genic region were calculated.

### Identification of drought-responsive sRNAs

A generalized linear model was fitted for each sRNA to identify drought responsive sRNAs. The response variable in the model is the read count of an sRNA, which were assumed to follow negative binomial distribution. The model contains two factors, DAW (day) and treatment, and their interactions. The DAW has eight factor levels (from 3 to 10) and the treatment has two factor levels (DS and WW). A deviance test of no interaction effect between DAW and treatment was conducted for each sRNA. The generalized linear model fit and test, assuming a negative binomial distribution for read counts, were implemented in DESeq2 (Love *et al*. 2014). sRNAs having at least five reads on average per sample were used for the statistical test, resulting in a p-value from each sRNA. A false discovery rate (FDR) approach was applied to account for multiple comparisons (Benjamini and Hochberg 1995). Significant sRNAs were declared using the 5% FDR as the cutoff. The script was deposited at GitHub (https://github.com/liu3zhenlab/sRNAs_drought).

### Clustering of drought-responsive sRNAs

Drought responsive sRNAs were subjected to clustering analysis using mclust (Fraley and Raftery 2007). For each drought-responsive sRNA, the Log2 of the ratio of the mean of DS expression (the normalized value) to the mean of WW expression (the normalized value) at a certain DAW (day) was determined, which represents the Log2 of the fold change in expression between DS and WW. Log2 ratio values were then used for the clustering analysis. The script was deposited at GitHub (https://github.com/liu3zhenlab/sRNAs_drought).

### Identification of significantly differentially expressed sRNAs between DS and water recovery

To test the null hypothesis that no difference in sRNA expression between two groups at 11 DAW, DR and water recovery (DWR), generalized linear model for the read count of each sRNA implemented in the DESeq2 package (version 1.4.5) was used (Love *et al*. 2014). A false discovery rate (FDR) approach was used to account for multiple tests (Benjamini and Hochberg 1995). The FDR 5% was used as the cutoff for declaration of differential expression.

### Enrichment analysis

The enrichment analyses were performed for determining if a certain type of category, such as a member of sRNA functional families, is over-represented in a selected group of sRNAs. To account for the biases read depth that influences the selection of members in a certain group, the resampling method in the GOSeq enrichment test (Young *et al*. 2010) with the bias factor of read depth, total reads across all the samples of a certain sRNA, was applied to enrichment analyses.

### Analysis of sRNA co-expression network

Drought-responsive sRNAs (FDR < 1%) were used to build co-expression sRNA network using Bioconductor package WGCNA (v1.51) (Langfelder and Horvath 2008). WW sRNA network was built using sRNA expression profiles in WW samples, and DS sRNA network was constructed using sRNA expression profiles in DS samples. The package WGCNA uses an appropriate soft-thresholding power to construct a weighted gene network. Modules of highly correlated sRNAs were identified using topological overlap measure (TOM) implemented in WGCNA. Module preservation analysis was also performed using WGCNA, with DS network as a test and WW network as a reference, and vice versa. An R script for network analysis has been deposited at GitHub (https://github.com/liu3zhenlab/sRNAs_drought).

### Identification of miRNAs

The database of mature miRNAs was downloaded from miRBase v22 (ftp://mirbase.org/pub/mirbase). In total, 325 mature B73 maize miRNAs from 174 miRNA genes were extracted. Any sRNAs discovered in this study identical to these mature miRNAs were annotated as known miRNAs.

ShortStack (v3.8.5) was used to *de novo* identify a set of miRNAs with the parameters (-- dicermin 18 --dicermax 30 --mismatches 0 --mincov 0.5rpm), and using B73Ref4 (version 4) as the reference genome (Axtell 2013b). ShortStack identified novel miRNA loci that did not overlap with any known miRNA genes. Any mature miRNAs from novel miRNA loci were referred to as novel miRNAs. Some mature miRNAs from ShortStack are not known miRNAs but from known miRNA genes. Combining both known mature miRNAs and all newly discovered mature miRNAs by ShortStack using our massive sRNA datasets, we updated the miRNA set, referred to as B73miRBase22plus.

### Identification of IsomiRs

IsomiRs are variants of the reference mature miRNAs (Morin *et al*. 2008). An isomiR in this study is a small RNA perfectly matching a pri-miRNA but with a different sequence from mature miRNAs in the B73miRBase22plus. Only 20-22 nt sRNAs identical to the plus-stranded sequence of a region of pri-miRNAs were referred to as isomiRs.

### Identification of ta-siRNAs

sRNAs matching ta-siRNA downloaded from tasiRNAdb (http://bioinfo.jit.edu.cn/tasiRNADatabase/) were defined as known tasiRNA. Also sequences of maize trans-acting siRNA 3 (TAS3) were retrieved from Dotto et al. (Dotto *et al*. 2014).

### Degradome analysis of drought-responsive miRNAs

Degradome raw reads were obtained from a previous maize miRNA study (Liu *et al*. 2014). After removing adaptor sequences and low-quality sequencing reads, clean reads were used to identify cleavage sites based on B73 cDNA sequences (5b+). CleaveLand 4.0 was implemented for degradome analysis with the default parameters (Addo-Quaye *et al*. 2009), which provides evidence for gene targeting by miRNAs or isomiRs.

### Prediction of miRNA targeted genes and GO enrichment analysis of targeted genes

psRNATarget (http://plantgrn.noble.org/psRNATarget/) was used to predict miRNA target genes (Dai and Zhao 2011). Gene targets of miRNAs were predicted based on B73 AGPv3.22 annotated transcript sequences with the expectation value no more than 1.5. Gene ontology (GO) enrichment of predicted miRNA-targeting genes was analyzed with AgriGO (Tian *et al*. 2017).

### Transposable element analysis of 24 nt genomic loci

sRNA genomic clusters, from the ShortStack result, predominant by 24-nt sRNAs were referred to as 24-nt genomic loci. RepearMasker (open-4.0.5) was used to identify sequences matching transposable elements with the maize transposon database. As a control, the “shuffle” module in the bedtools was employed to randomly select intervals simulating the number and sizes of genomic intervals of 24-nt loci.

### Data Availability

The datasets supporting the conclusions of this article are included within the article and its supplemental materials. Supplemental files available at FigShare. All sRNA sequencing raw data were deposited at Sequence Read Archive (SRA) (accession number: SRP081275).

## Results

### Physiological changes of seedlings under drought conditions

Maize seedlings were subjected to drought over a period of nine days (Figure 1A). Three-day-old B73 seedlings after germination were subjected to two treatments, drought stress (DS) and well-watered (WW). Above ground tissues (referred to here as leaves) were collected at 3 to10 <u>D</u>ays <u>A</u>fter <u>W</u>ithholding water (DAW) or with watering with two biological replicates at each day. At 10 DAW, some seedlings from the DS treatment group were subjected to two treatments: continuously withholding water (DS) and re-watering, both of which were sampled at the 11th day. Two biological replicates were collected, resulting in two additional DS samples on day 11 and two re-watering samples at one day after addition of water at day 10. A total of 36 plant samples were processed. Compared to WW seedlings, DS-treated seedlings showed severe stressed phenotype by 8 DAW. Soil water content (SWC) decreased in the DS treatment from ~60% to 20% in the same period (Figure 1B). Leaf relative water content (RWC) of DS seedlings also decreased upon the drought treatment, at a low declining rate from 3 to 7 DAW and a high rate after 7 DAW (Figure 1C). Leaf relative electrical conductivity (REC), which is a measure of cellular damage, exhibited strongest response to drought between 8 and 9 DAW (Figure 1D), indicating that leaf cells began to experience damage after 8 DAW under drought conditions. The DS-treated seedlings showed visible stressed phenotypes after 10 DAW. When re-watered at 10 DAW, the DS-treated plant seedlings were visibly recovered at 11 DAW.

**Figure 1.**
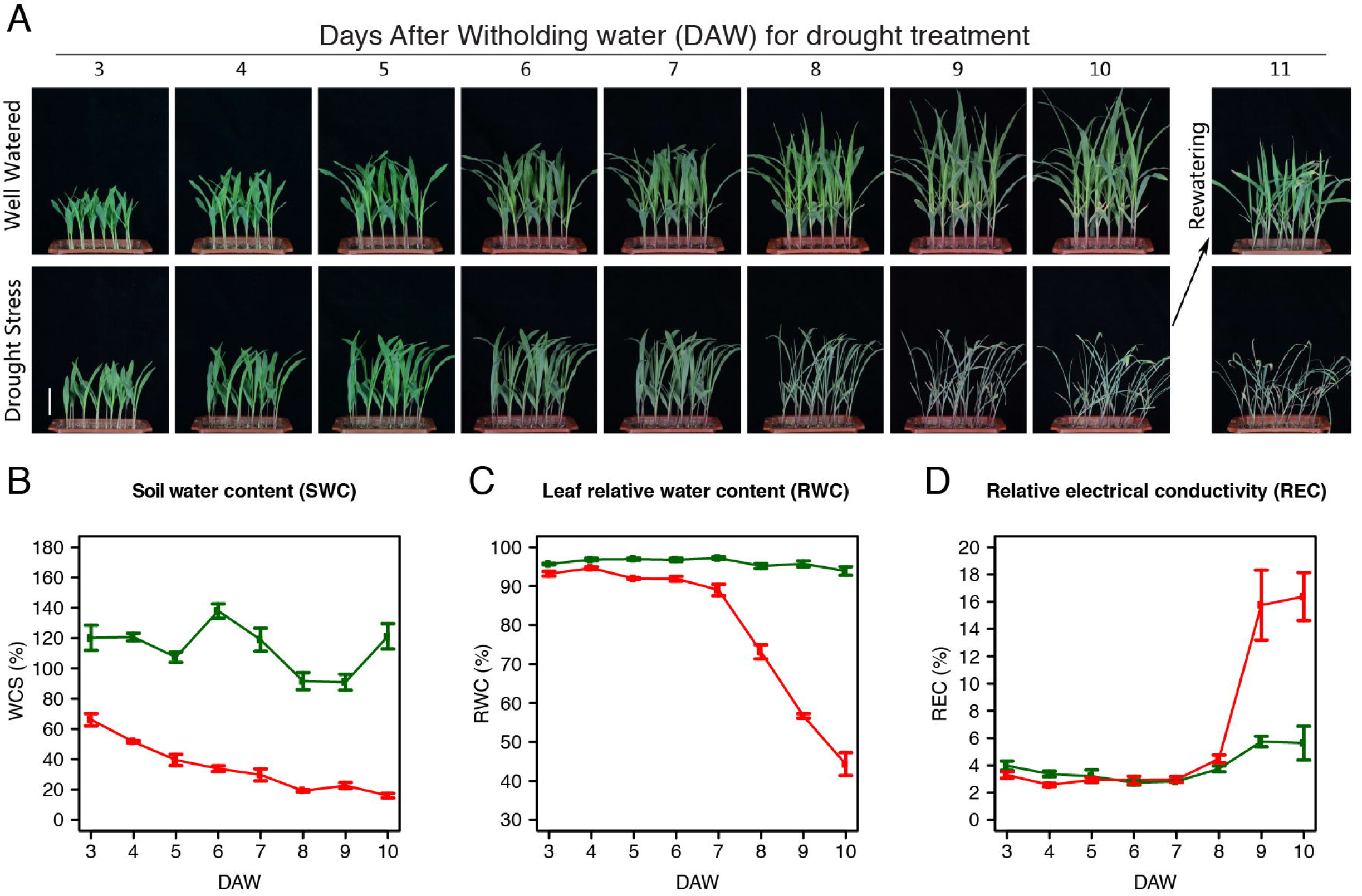
Morphological and physiological changes of maize seedlings during drought stress. (A) Three-day-old B73 seedlings were subjected to gradual drought stress or well-watered conditions. The photos were taken at each day from 3 to 11 days. Bar=5cm. (B) The changing curves of water content of soil (SWC) from five replicated pots of each data point. (C) Leaf relative water content (RWC) of seedlings along days. (D) Leaf relative electrical conductivity (REC) of seedlings along days. Red and green curves represent plants under drought stress and well water, respectively. Five seedlings were pooled as one replicate, and five independent biological replicates were conducted to determine RWC and REC. Vertical lines represent standard errors.

### Characterization of sRNAs

The 36 RNA samples were extracted for sRNA sequencing, resulting in more than 886.6 millions of 50 bp single-end reads, from 20.5 to 34.2 millions reads per sample. On average, 97.5% of reads were retained after adaptor and quality trimming of each sample (**Table S1**). The majority of sRNAs were between 18 and 26 nucleotides (nt). The 24-nt sRNA length class was the largest, followed by the 21 nt and 22 nt sRNA classes (Figure 2A). The same pattern of length distribution was observed across all the samples, indicating that the drought treatment did not alter the global pattern of sRNA lengths.

**Figure 2.**
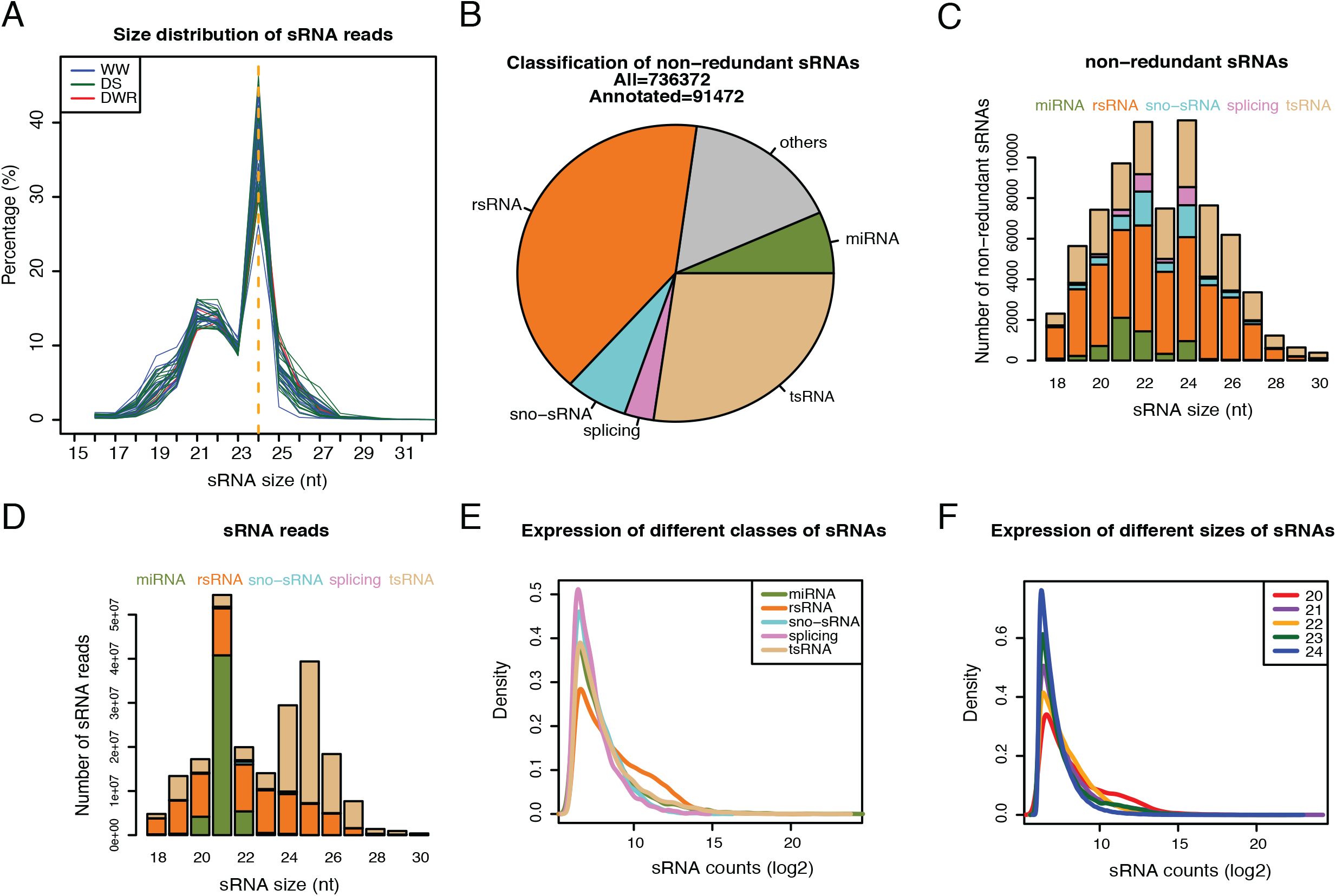
Characterization of sRNAs. (A) Proportions of sRNAs of different lengths in all samples. Each curve represents a sample. WW, DS, and DWR, represent well-watered, drought stress, and drought water recovered plants, respectively. (B-D) Overview of genomic copy number, lengths, functional categories, and expression of NR-sRNAs from all the samples. (B) Pie chart of distribution of different classes of sRNAs. Others represent sRNAs that were not unambiguously categorized. (C) Stacked barplot of different functional classes of NR-sRNAs at varying sizes of sRNAs from 18 to 30 nt. (D) Stacked barplot of different functional classes of sRNA reads, representing expression levels, at varying lengths of sRNAs from 18 to 30 nt. (E) Density plots of expression levels of different functional classes of sRNAs. Density on the y-axis represents the probability of sRNA occurrences. (F) Density plots of expression levels of different lengths of sRNAs. Density on the y-axis represents the probability of sRNA occurrences.

All sRNA reads from the 36 samples were merged, and sRNAs with at least 72 reads were retained. Removing redundant reads with the same sequence for each sRNA resulted in a non-redundant sRNA (NR-sRNAs) set of unique 736,372 sRNAs (134,283 NR-sRNAs used in the later time-series statistical analysis were listed in **Table S2** and **Table S3**. The NR-sRNAs set was annotated using the Rfam database (Rfam11.0). 12.4% (91,473) of the NR-sRNAs could be unambiguously annotated with regard to function (see Methods). Among the Rfam-annotated subset of sRNAs, rsRNAs, tsRNAs, and miRNAs are the most abundant, comprising of 40%, 27%, and 7%, respectively (Figure 2B). The rsRNA and tsRNAs represented nearly 70% of all annotated NR-sRNAs, while miRNAs distributed in a slightly narrower length range of 18 to 24 nt and a peak length at 21 nt (Figure 2C). Of 21 nt NR-sRNAs, 22% are miRNAs from approximately 65% of the total 21-nt sRNA reads (redundant sRNAs), indicating that some 21 nt miRNAs were highly expressed (Figure 2C, 2D). Indeed, the single sRNA showing the highest abundance is a miR159, with 14.8 million reads.

### Genome organization of NR-sRNAs in B73

The copy number of individual NR-sRNAs in the B73 genome was estimated by both mapping reads to the B73 reference genome (reference-based) and analyzing sequences present in whole-genome-shotgun sequence reads (WGS-based) (see Methods). A small number of NR-sRNAs, 20,452, were excluded based on alignment to either chloroplast or mitochondria DNA. Among the remaining NR-sRNAs (N=705,920), perfect matches for 93.2% of the NR-sRNAs were identified in either the B73 reference genome or the B73 WGS data. The absence of perfect matches for 6.8% of the NR-sRNAs was attributed to incomplete B73 genome assembly, contamination, sequencing errors, and/or RNA editing (Liang and Landweber 2007; Schnable *et al*. 2009). The estimations of copy number from the two approaches were largely consistent (**Figure S1**). Both estimations indicated that most NR-sRNAs are from low-copy genomic loci (1-2 copies) except for NR-sRNAs from rRNA and tRNA (**Figure S2**). NR-sRNAs of differing lengths exhibit varying mixtures of low- and high-copy loci (**Figure S3**). The 24 nt sRNAs are mostly single copy in the genome, while a high proportion of 21-23 nt sRNAs are derived from either low-copy or very-high-copy genomic loci. Outside of the 21-24 sRNA range, NR-RNAs from highly repetitive genomic regions are dominant (**Figure S3**).

A linear association between expression level and genomic copy number of sRNAs was not observed (**Figure S4**). Genomic single-copy NR-sRNAs can be highly expressed. For example, the single copy miR168 locus was expressed at a high level (138,292 reads). Conversely, the expression of most genomic high-copy NR-sRNAs was low. Some high-copy NR-sRNAs were highly expressed, such as rsRNAs. Analysis of sRNA expression profiles based on functional classes also showed that high proportions of splicing sn-sRNAs and sno-sRNAs exhibit low expression, while many rsRNAs were expressed at a high level (Figure 2E). The 23 and 24 nt sRNAs, regardless of functional classes, were mostly expressed at a low level, while 20-22 nt sRNAs tended to be expressed at relatively higher levels (Figure 2F). Compared to 21-24 nt sRNAs as a whole, a higher proportion of 20 nt sRNAs were highly expressed (Figure 2F).

### Distinct histone modifications at genomic regions of different classes of sRNAs

Histone modification status at genomic regions of sRNAs was collected from genome-scale data repositories for B73 seedlings, including multiple histone modifications: H3K27me3, H3K36me3, H3K4me3, H3K9ac, and H3K9me2. The lack of biological replication and low depth of most chromatin modification data limited assessment of histone modification levels for each sRNA locus. Therefore, the mean of histone modification levels of genomic regions in each functional sRNA class was used to represent the overall genomic modification level of each sRNA functional class. To avoid systematic biases, we compared histone modifications among different functional classes at the same sRNA length (Figure 3, **Figures S5-S8** and **Table S4**). Average histone modification levels on different functional classes showed that both miRNAs and sno-sRNAs in size of 20, 21, and 23 nt were predominately found in open chromatin regions, which were characterized by high modification levels of two hallmarks of open chromatin regions, H3K4me3 and H3K9ac. The H3K4me3 signal at sno-sRNA genomic regions was much higher than those of genomic regions of any other functional sRNA classes across all lengths from 20 to 24 nt. Genomic regions of 21 nt miRNAs and sno-sRNAs, overall, had moderate levels of a silent chromatin mark H3K9me2, and genomic regions of 20 and 23 nt miRNAs and sno-sRNAs exhibited low levels of H3K9me2 signals. H3K9me2 is associated with CHG (where H is A, T, or C) cytosine methylation (Stroud *et al*. 2013; West *et al*. 2014), indicating that genomic regions producing miRNAs and sno-sRNAs, on average, exhibited low CHG cytosine methylation. At the lengths of 20, 21, 23 nt, miRNA genomic regions had high levels of H3K27me3, a repressive chromatin mark associated with gene silence (Gan *et al*. 2015), and relatively low H3K36me3 levels that are generally positively associated with transcriptional activity but also were found to be enriched at heterochromatin regions (Chantalat *et al*. 2011). The sno-sRNA genomic regions, relative to miRNA genomic regions, exhibited the opposite modification pattern with low H3K27me3 levels and high H3K36me3 levels. Genomic regions of rsRNAs, splicing sn-sRNAs, and tsRNAs had similar histone modification patterns, namely, low levels of H3K27me3, H3K36me3, H3K4me3 and H3K9ac and a high level of H3K9me2 across all lengths from 20 to 24 nt.

**Figure 3.**
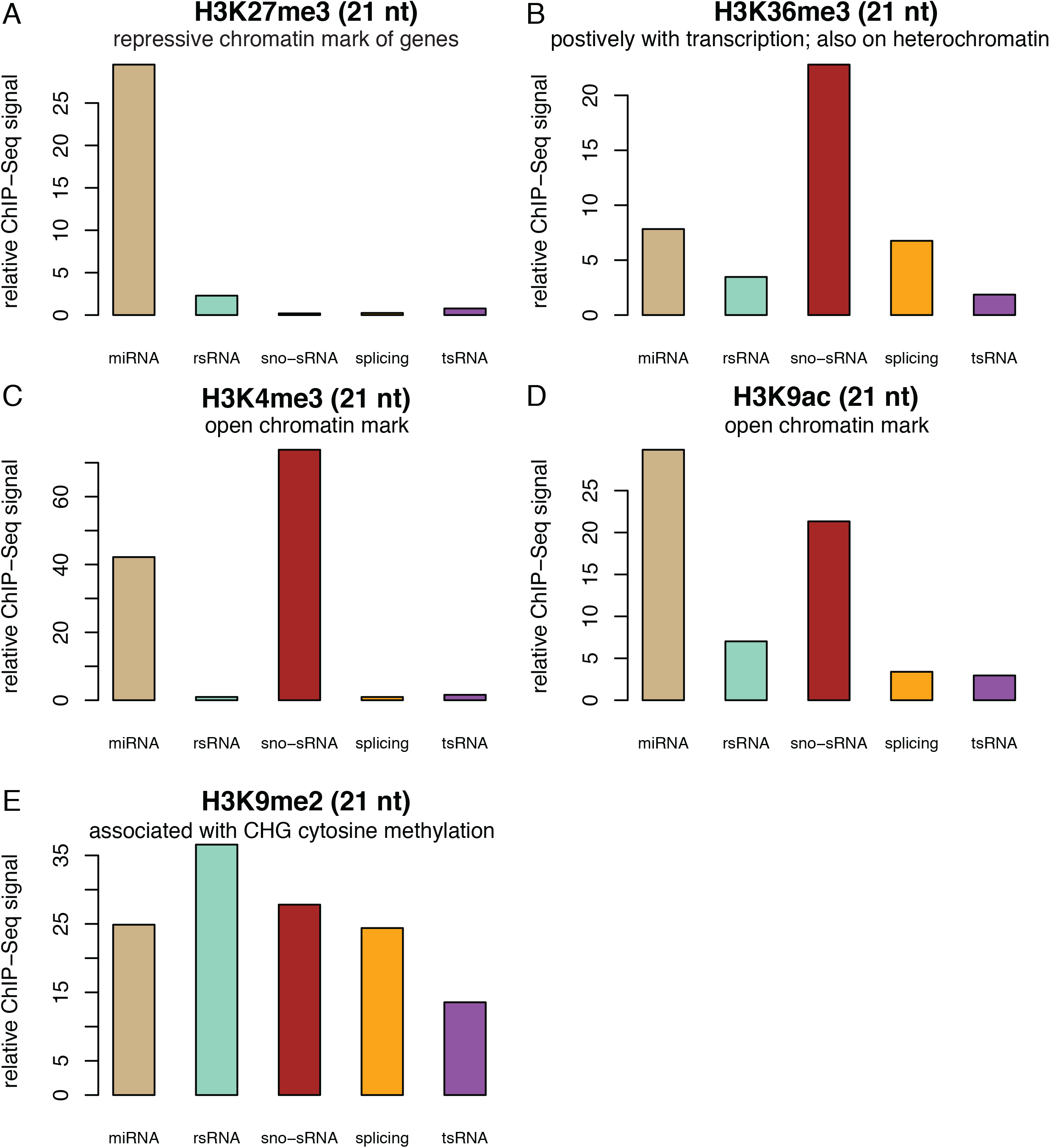
Modification levels of five epimarks on sRNA genomic regions. The average ChIP-Seq signals, represented by read depths of ChIP-Seq, of five epimarks were determined and normalized by sequencing library sizes separately. Heights of bars represent relative histone modification levels. The general function of each epimark is briefly described in the subtitle of each barplot.

Through converting all lengths of sRNAs to the same 18 nt length, average signals of histone modifications were compared among genomic regions producing different lengths of sRNAs. As the control, 18 nt DNA fragments were also randomly sampled from different genic regions, and their mean signals of histone modifications were determined. The results showed that, except for H3K36me3, all epimarks shared a similar trend that the modification signals were at a relatively high level for genomic regions of small lengths of sRNAs and gradually decrease until at 24 or 22 nt, followed by elevated modification signals (**Figure S9**). For open chromatin marks H3K4me3 and H3K9ac, on average sRNAs exhibited lower levels relative to promoters and first exons but similar levels to internal exons, introns and last exons. Our result showed that H3K9me2 was generally at much higher levels on sRNA genomic regions relative to genic regions, of which 24 nt sRNAs whose genomic regions had the closest H3K9me2 signal to genic regions. That indicated that the CHG cytosine DNA methylation at genomic regions of 24 nt sRNAs is generally low.

### Identification of drought-responsive sRNAs

A statistical test was performed to detect any interaction between drought stressed and well-watered plants for each sRNA that had a minimum five sRNA reads per sample over the 3 to 10 DAW period. The analysis revealed that 6,646 of the total 134,283 sRNAs exhibited interactions between the DAW and the treatments at the 5% false discovery rate (FDR) level (**Table S3**). Interacting sRNAs showing different responses under DS and WW conditions at certain DAWs were scored as drought-responsive sRNAs. The rsRNAs and 22 nt sRNAs are the two predominant groups in the drought-responsive sRNA set (**Figure S10**). The DS-to-WW ratios of sRNA expression were further subjected to cluster analysis using mclust (Fraley and Raftery 2007), resulting in 10 clusters. The sRNAs of clusters 3, 4, 5, 7, and 9 exhibited a pattern of up-regulation under drought stress (Figure 4A-F), while sRNAs of clusters 1 and 8 showed a pattern for down-regulation (Figure 4G-I). More than five times up-regulated sRNAs (N=4,373) were detected than down-regulated sRNAs (N=816) under drought stress (Figure 4). The enrichment analyses indicate that rsRNAs and splicing sn-sRNAs were over-represented in up-regulated sRNAs, while miRNAs and sno-sRNAs were over-represented in down-regulated sRNAs. Additionally, sRNAs of clusters 2 and 6 exhibited transiently down-regulation on drought (transiently down-regulation group, N=1,325), which were down-regulated at around 7 DAW when drought stress became intense, followed by a gradual recovery of expression (Figure 4J-L). The enrichment analysis indicates that miRNAs and sno-sRNAs are significantly over-represented in transiently down-regulated sRNAs.

**Figure 4.**
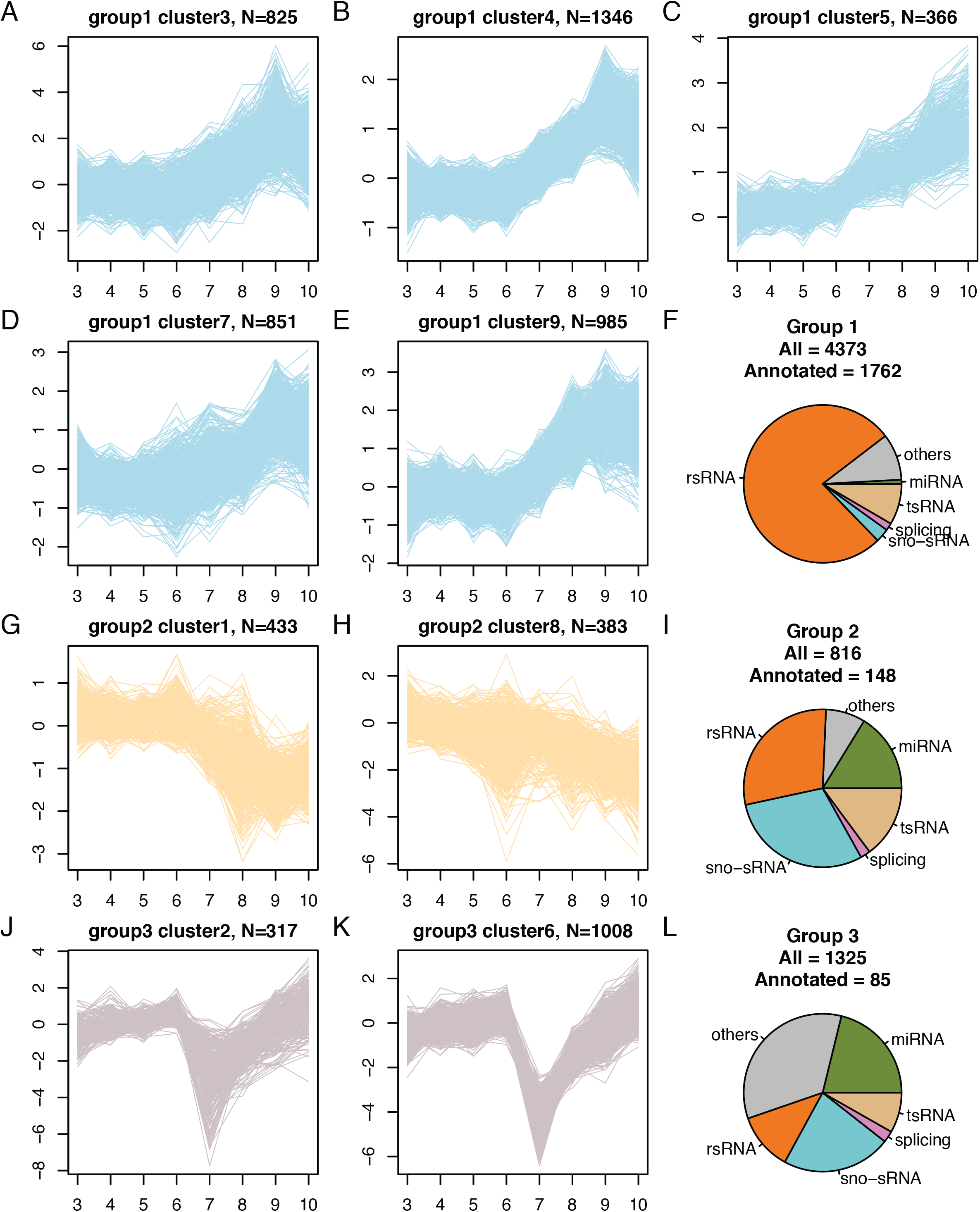
Major clusters of drought-responsive sRNAs. Drought-responsive sRNAs were subjected to clustering using the software mclust, which ended with 10 clusters. Nine major clusters (A-E, G, H, J, K) were classified into three groups, up-regulated (light blue), down-regulated (light orange), and transiently down-regulated (light purple). Each curve represents an average sRNA expression ratio of drought stress to well-watered with a Log2 transformation from two biological replicates along DAW. Three pie charts designate proportions of different classes of sRNAs that were functionally annotated in each of the three clustering groups: up-regulated (F); down-regulated (I); transiently down-regulated (L).

A comparison of sRNA expression was performed between two additional seedling groups at 11 DAW, DS and drought water recovery (DWR), which was re-watered on 10 DAW. Using the 5% FDR cutoff, 7,140 sRNAs were differentially expressed between two groups, of which 2,264 and 4,876 sRNAs were up-regulated and down-regulated in DWR relative to DS, respectively, and 486 were identified as drought-responsive sRNAs in the time-series analysis (**Table S3, S5**). The 473 sRNAs (out of 486) were classified into three groups in the time-series analysis: Down-regulated (N=43), up-regulated (N=426), and transiently down-regulated (N=4). All 43 sRNAs from the down-regulated group were up-regulated after DWR. Of 426 sRNAs in the up-regulated group, 76.3% (325/426) sRNAs showed decreased expression in DWR, while 23.7% (101/426) were continuously up-regulated even with water recovery. All four sRNAs in the transiently down-regulated response group were up-regulated after re-watering. Overall, the expression levels of most drought-responsive sRNAs were restored towards levels of well-watered plants upon re-watering.

### Characteristics of co-expression networks of drought-responsive sRNAs

DS and WW weighted co-expression networks were constructed using WGCNA (Langfelder and Horvath 2008). Both networks consist of a subset of drought-responsive sRNAs with the FDR cutoff of less than 1% from the drought response statistical test. The DS and WW networks were built using normalized sRNA counts of DS and WW samples, respectively (Figure 5A, 5B, **Table S3**). Network statistics indicate intrinsic differences between the two networks (**Table S6**). Although the DS and WW networks share similar network clustering coefficients, network centralizations, and network densities, the DS network (Figure 5B) has the smaller network diameter and lower heterogeneity, indicating expression of these drought-responsive sRNAs were more correlated upon drought stress or tended to be co-expressed in response to drought stress.

**Figure 5.**
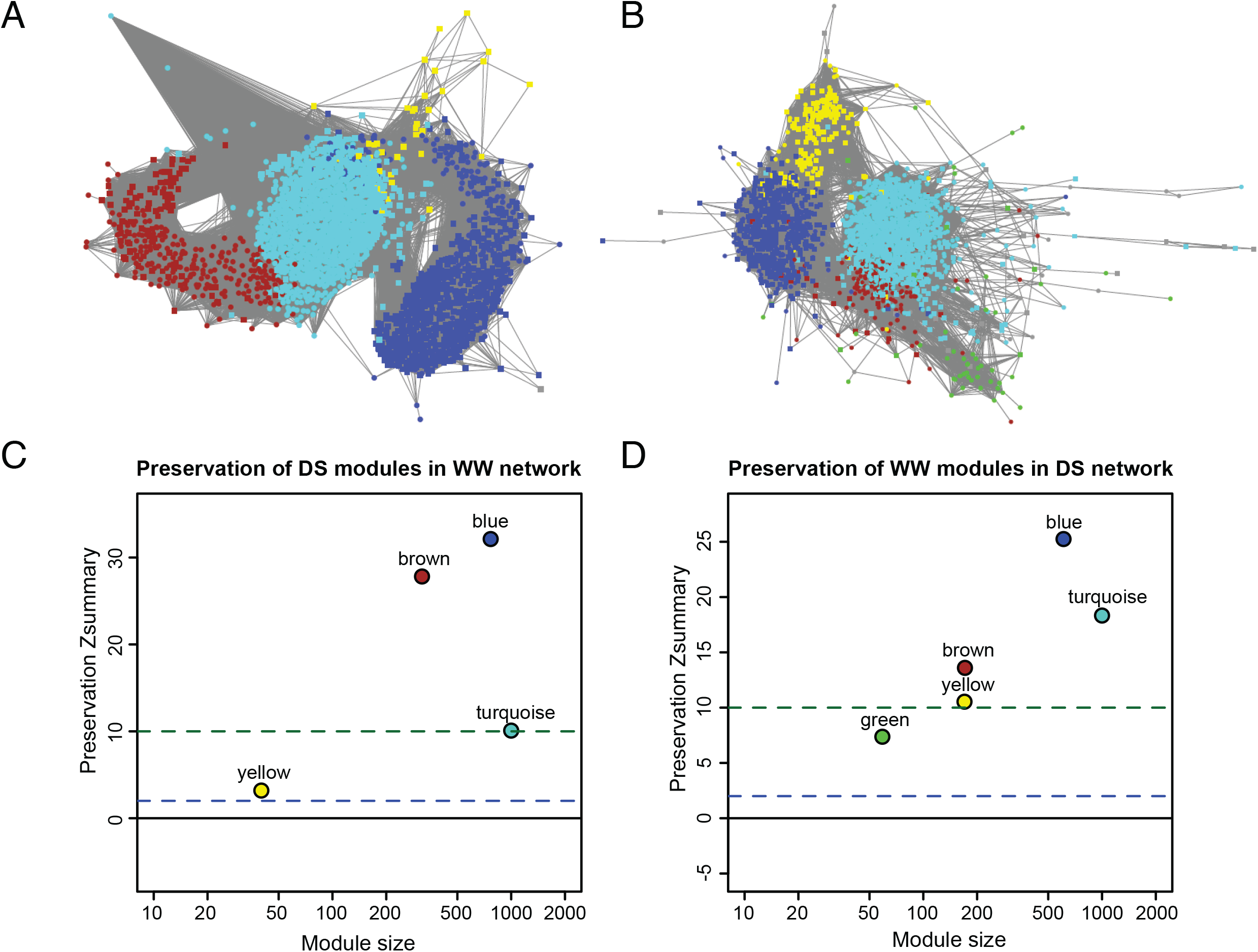
sRNA co-expression networks (A) Visualization of the DS network using Cytoscape, each node represents an sRNA and each line is the edge connecting sRNA nodes. Five modules (sub-network) were highlighted by different colors. (B) Visualization of the WW network. Six modules (sub-networks) were highlighted by different colors. Note that assignment of colors in (A) and (B) are sort of independent. The same color might not represent the same group of sRNAs. (C) Result of the module preservation analysis performed to evaluate whether a module identified in the DS network is preserved in the WW network. The color code corresponds to that used in (A). (D) Result of the module preservation analysis of the WW network in comparison with the WW network. The color code corresponds to that used in (B).

Modularity analysis in the DS network and the WW network further revealed that the two networks have different topology structures. Modularity analysis included two steps: module identification and module preservation analysis. Modules are sub-networks, consisting of co-expressed sRNAs. The sRNAs in the same module are similar in expression to some degree, thereby are likely associated each other. Module preservation analysis is used to determine if the topology of a network module identified in one network changes in the other network. For example, a module is considered preserved in the DS network, if its topology, based on preservation statistics, largely remains in the WW network. The module preservation analysis identified a preserved module (blue module) in the DS network compared to the WW networks (Figure 5C) and a preserved module (blue module) in the WW network in comparison to the DS networks (Figure 5D). Most sRNAs (N=546) in two blue modules overlapped, of which more than 95% are from the transiently down-regulated group (**Table S3**). The result indicated transiently down-regulated sRNAs tended to be co-regulated in both drought and well-watered conditions. On the other hand, these sRNAs exhibited a transient down-regulation to drought, which might serve as the signal to induce downstream drought responses. Of 546 overlapping sRNAs, 343 and 178 are 22 nt and 24 nt sRNA, respectively, and a few were functional annotated with the Rfam database (6 miRNAs and 9 sno-sRNAs). The module preservation analysis also revealed differences between modules in the DS and WW networks. The yellow module in the DS network is the least preserved module, indicating sRNAs of the module were perturbed in response to drought stress (Figure 5C). Indeed, the yellow module consists of 38 sRNAs that were down-regulated upon drought stress. In the WW network, the green module is the least preserved one, and most sRNAs were up-regulated upon drought.

### Identification of drought-responsive miRNAs and the corresponding targeted genes

sRNA homologous to Rfam miRNAs were referred to as miRNAs hereinbefore. We refined the miRNA set based on the dedicated miRNA database, miRBase (Kozomara and Griffiths-Jones 2014), and *de novo* discovery of miRNAs from our massive datasets. We employed the ShortStack pipeline (Axtell 2013b) and identified 53 miRNA loci of which 47 loci are known maize miRNA genes in miRBase (v22) containing 174 miRNA genes. We found 59 new mature miRNAs from, including 47 mature miRNAs from known miRNA loci but with different sequences of mature miRNAs, as well as 12 mature miRNAs from 6 novel miRNA loci. Requiring at least an 18 nt match with at least 90% identity, homologs of miRNAs from three novel miRNA loci (Cluster_23765, Cluster_27697, and Cluster_45700) were identified in MIR1878, MIR156c, and MIR166d, respectively. We combined both known and newly discovered mature miRNAs to create a new miRNA set referred to as B73miRBase22plus (**Table S7**) that contains 180 miRNA genes producing 392 mature miRNAs, of which 244 are non-redundant miRNAs (**Table S8**). We also identified 608 isomiRs that are in length of 20-22 nt and identical to a region of a pri-miRNA sequence, but different from 392 mature miRNAs in sequence (**Table S9**).

Some miRNAs were highly expressed. The top eight most highly expressed miRNAs belong to six families: miR159, miR168, miR396, miR156, miR169, and miR167 (**Table S8**). Although highly expressed miRNAs, statistically, are most likely to be detected, none of the top 25 miRNAs showed evidence of regulation under drought condition, indicating that expression levels of most highly expressed miRNAs were kept at relatively stable levels under drought stress. In total, 21/244 miRNAs and 18/608 isomiRs showed significantly drought responses (Table 1). Most drought-responsive miRNAs (N=13) were down-regulated by drought treatment, while four were up-regulated. The remaining four were not categorized to any of the three major cluster groups. The 21 drought-responsive miRNAs belong to 13 families, including miR1432, miR156, miR164, miR166, miR167, miR168, miR171, miR319, miR390, miR398, miR399, miR408, and miR528 (Table 1). The miR390a-3p or miR390b-3p (miR390a/b-3p) of the miR390 family was drought responsive. But no significant regulation on drought was observed for miR390a/b-5p (AAGCUCAGGAGGGAUAGCGCC) that cleaves trans-acting siRNA 3 (TAS3) loci to produce ta-siRNAs (Allen *et al*. 2005; Williams *et al*. 2005; Dotto *et al*. 2014; Xia *et al*. 2017). Predicted TAS3 ta-siRNAs triggered by miR390a/b-5p were either low expressed or with no significant regulation upon drought stress (**Table S10**). For isomiRs, 7, 8, and 3 were in down-regulation, up-regulation, and uncategorized groups, respectively, adding two additional miRNA families, miR396 and miR444, showing drought responses. Notably, multiple isomiRs, and mirR156i-3p, from the miR156 family were up-regulated on drought (Table 1). However, miR156j-3p was down-regulated, implying that family members play divergent regulatory roles.

**Table 1.**
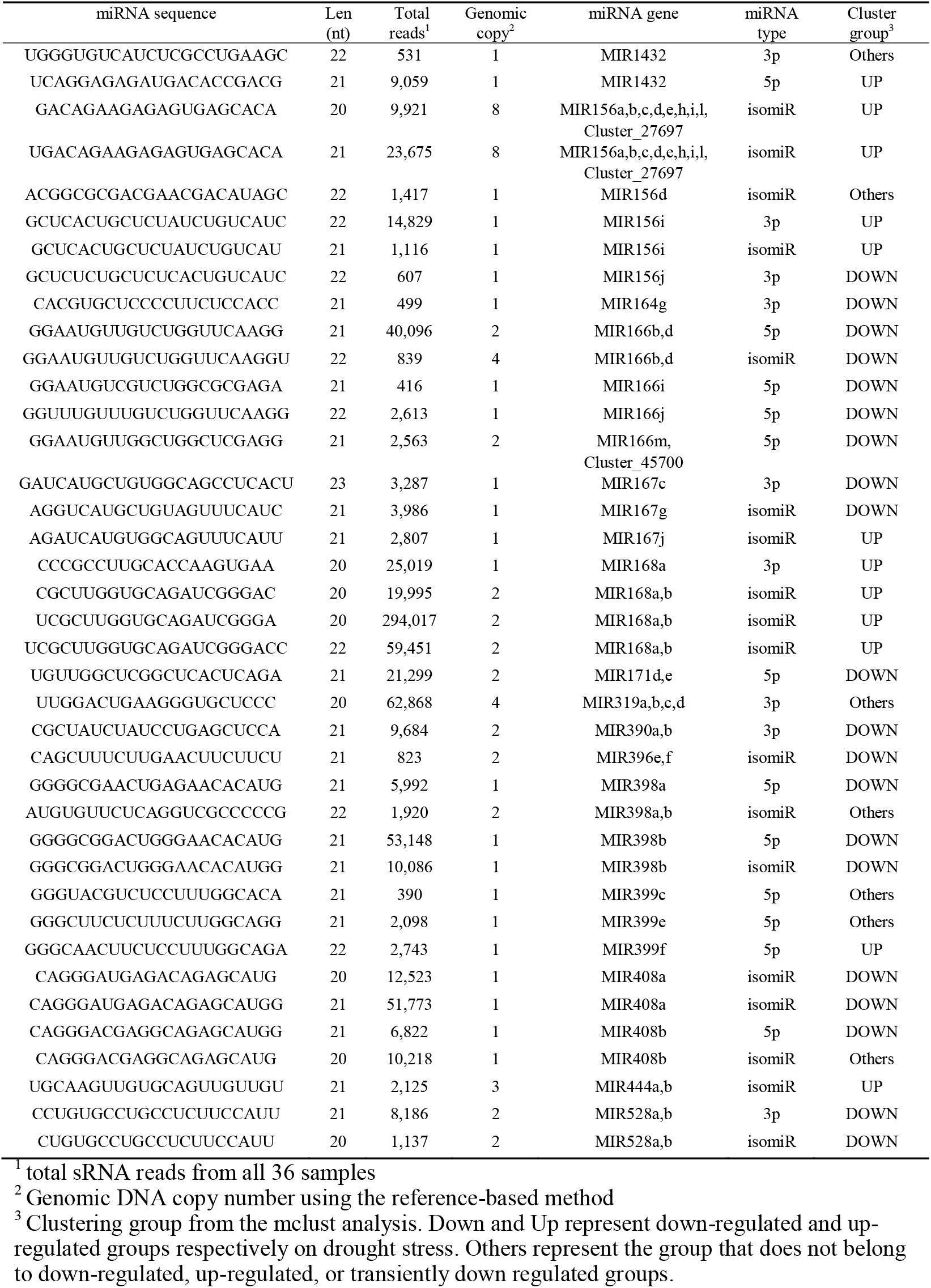
The list of drought-responsive miRNAs

Targeted protein-coding genes of 21 miRNAs and 18 isomiRs responded to drought were predicted with the psRNATarget tool (Dai and Zhao 2011). In total, 67 pairs of gene-miRNA, including 43 non-redundant genes, were predicted to be targeted by 18 drought-responsive miRNAs and isomiRs (**Table S11**). GO enrichment analysis showed that 43 miRNA-targeting genes are highly enriched in DNA binding function (GO:0003677, p-value = 2.1E-16) and nucleus cell component (GO:0005634, p-value = 6.1E-16) (**Table S12**), suggestive of considerable impacts of miRNAs on the genes regulating transcription under drought stress. Nearly half of targets (18/43) are putative SPL (Squamosa promoter binding protein-like) transcription factors, and 17/18 are targeted by two isomiRs of the miR156 (GACAGAAGAGAGUGAGCACA and UGACAGAAGAGAGUGAGCACA). SPL genes have been reported to be associated with miR156 under drought condition in multiple plant species, such as rice (Nigam *et al*. 2015) cotton (Wang *et al*. 2013), alfalfa (Arshad *et al*. 2017), and maize (Mao *et al*. 2016). In our result, both SPL-targeting miR156 were up-regulated upon drought (Figure 6), indicating the possible regulation in expression of SPL genes through miRNAs during drought treatment. Another drought-responsive miRNA miR319a/b-3p (UUGGACUGAAGGGUGCUCCC) was predicted to target one MYB and two TCP transcription factors (GRMZM2G028054, GRMZM2G089361, GRMZM2G115516) (Zhang *et al*. 2009; Liu *et al*. 2014). This miR319a/b-3p remained at a low expression level under high drought stress (**Figure S11**). Presumably, the expression of targeted genes was under a low level of suppression imposed by miR319 under drought condition. Indeed, one of three genes GRMZM2G115516 was up-regulated >4 times on drought (**Table S11**) (Liu *et al*. 2015). The transcriptional regulation of genes targeted by isomiRs of miR156 and miR319a/b-3p was well supported from degradome sequencing data (**Table S11**), which were used to identify miRNA cleavage sites (Shen *et al*. 2013; Zhai *et al*. 2013; Liu *et al*. 2014).

**Figure 6.**
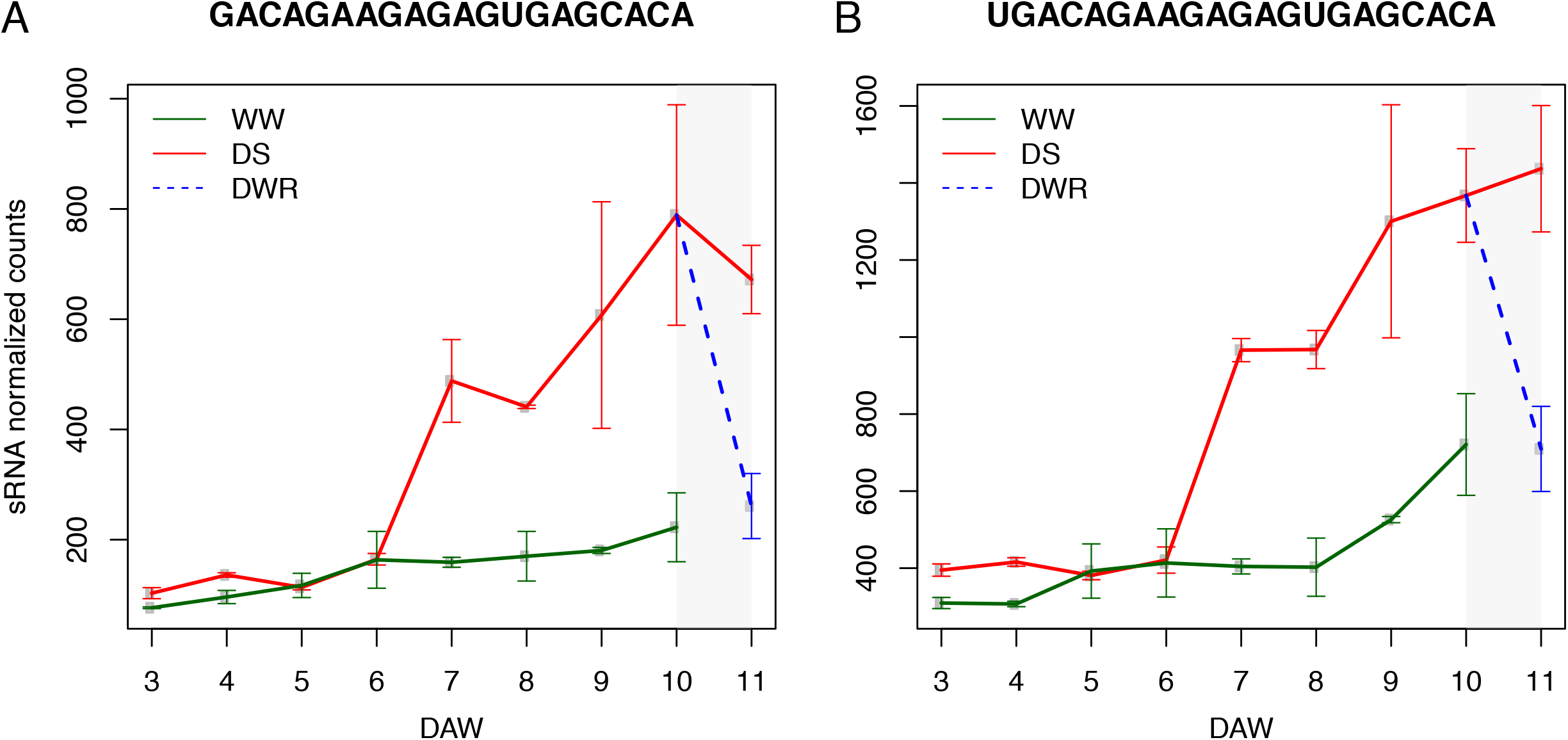
Time-series expression profiles of two miR156 targeting SPL genes (A, B) Normalized counts of each miR156 (y-axis) were plotted along 3-11 DAW. WW, DS, DWR represent well-watered, drought stress, drought water recovery, respectively. A sequence on the top of each plot is the miR156 sequence. Each error bar represents the range of a standard error above and below each mean value.

## Discussion

In this study, sRNA sequencing was performed on samples of maize seedlings under drought stress (DS) and well-watered (WW) conditions. The sRNAs were characterized with respect to sRNA lengths, functional class, as well as copy number and epigenetic modifications of sRNA genomic regions. Genomic copy number analysis indicates that most 18-20 nt and 25-30 nt NR-sRNAs and approximately half of the 21-23 nt NR-sRNAs are derived from high-copy genomic repeats. The 24 nt sRNAs were the predominate species among single-copy sRNAs in this study, which is inconsistent with the observations in most other plant species. In fact, 24 nt sRNAs are generally referred to as heterochromatic siRNAs and are primarily derived from intergenic and/or repetitive genomic regions (Dunoyer *et al*. 2007; Kasschau *et al*. 2007; Axtell 2013a). However, 24 nt sRNAs were also recently shown to be enriched in euchromatic regions with low DNA cytosine methylation in an independent maize study (He *et al*. 2013), which is consistent with our observation. Based on ShortStack sRNA genomic mapping, 24-nt sRNA genomic loci were largely located at intergenic regions, but closer to protein-coding genes compared to randomly shuffled simulated loci (**Figure S12**). The proximity of 24-nt sRNA genomic loci to protein-coding genes, particularly highly expressed genes, was previously observed (Lunardon *et al*. 2016), and the 24-nt sRNA was proposed to function to reinforce silencing of transposable elements close to active genes (Li *et al*. 2015a). Our transposon analysis found that 24-nt sRNA genomic loci were overrepresented at regions containing DNA transposon elements but under-represented at regions containing LTR retrotransposon elements, *Copia* and *Gypsy*’(**Table S13**), suggesting the 24-nt sRNA might be more critical for silencing of DNA transposon elements. Compared to other lengths of sRNAs, genomic regions generating 24 nt sRNAs exhibited low histone modification levels for all histone epimarks examined. Given that most 24 nt sRNAs are generated by PolIV, heavy nucleosome loading and/or strong histone modifications of examined epimarks are likely not prerequisites for transcription via PolIV (Li *et al*. 2015b; Lunardon *et al*. 2016).

High proportions of sRNAs with two genomic copies were found in 21 and 22 nt sRNAs but not in 23 or 24 nt sRNAs. Production of most 23 and 24 nt sRNAs requires RNA-dependent RNA polymerase 2 (RDR2) to form dsRNAs and do not require multiple genome copies for optimal function (Nobuta *et al*. 2008). Two identical copies in the genome could increase the chance for the expression of sense and antisense transcripts to form NAT-siRNAs, which indicates that many 21 and 22 nt sRNAs might be NAT-siRNAs. Genomic regions of 21 and 22 nt sRNAs have the highest modification levels of H3K36me3 among all lengths of sRNAs, resembling H3K36me3 modification levels of internal genic regions (internal exons and introns). The high H3K36me3 signals of 21 and 22 nt sRNAs genomic regions are likely contributed by sno-sRNA genomic regions, which exhibited the highest H3K36me3 modification levels. Our results revealed the complexity of histone modifications of plant sRNA genomic regions. However, the lack of high depth of epimark data as well as the different experimental sources between epimark information and sRNA expression data restrict the conclusion about their correlation at a single locus level. Future stratification based on sRNA length, function, as well as genomic and more informative epimark information of sRNA genomic regions would be useful for understanding biogenesis and cellular function as well as further classification of sRNAs.

Characterization of drought-responsive sRNAs indicates that sRNAs are differentially expressed in response to drought stress. The miRNAs of maize were clustered into three groups based on expression patterns, namely, up-regulated, down-regulated, and transiently down-regulated upon drought stress and over-represented in the down-regulated group, in which miRNAs were approximately 4.8x enriched. The miRNAs and cognate gene targets are involved in drought stress responses in many plant species such as *Arabidopsis* (Butler *et al*. 2008), rice (Zhou *et al*. 2010; Fang *et al*. 2014), soybean (Axtell 2013b) and poplar (Shuai *et al*. 2013). Drought-induced miRNAs suppress their target mRNAs, while down-regulated miRNAs result in the de-repression of the target mRNAs (Ferdous *et al*. 2015). The miRNAs may exhibit distinct responses to drought stress in different plant species (Zhai *et al*. 2015). For example, miR168a/b down-regulated on drought in rice (Zhou *et al*. 2010), but was induced in response to drought stress in maize. We have identified 39 drought-responsive miRNAs or isomiRs, as well as their potential gene targets. Detailed studies on their regulatory networks and their functional divergence among species or genotypes within a species would be valuable to modulate miRNA-mediated pathways for improving drought tolerance of plants.

In addition to miRNAs, sRNAs derived from rRNAs, tRNAs, snoRNAs, and splicing snRNAs were also differentially regulated under drought condition. rRNAs are an essential component of ribosomes and catalyzes protein assembly. rsRNAs (small RNAs derived from rRNAs) were over-represented in the up-regulated sRNA group. rsRNAs were significantly enriched in down-regulated sRNAs after addition of water at 10 DAW. Thus, drought response involves an increase of rsRNAs, which is, in turn, suppressed when water was supplied. Transfer RNAs (tRNAs) play an essential role in protein synthesis. Although tsRNAs (small RNAs derived from tRNAs) are not enriched in either up- or down-regulated sRNAs groups, up-regulated tsRNAs are almost seven times more represented than down-regulated tsRNAs (148/22), which is higher than the ratio of all up-regulated sRNAs to all down-regulated sRNAs (4,373/816). A barley sRNA study also found that tsRNAs, overall, have a tendency to be up-regulated under drought condition (Hackenberg *et al*. 2015). Both rsRNA and tsRNA were abundant at all lengths from 18 to 27 nt, implying that the cleavage activity of rRNA and tRNA is not size-specific. Likely an unknown RNase III member is involved in rRNA or tRNA cleavage, producing sRNAs with a broad range of lengths (Wu *et al*. 2000). Splicing sn-sRNAs, derived from splicing snRNAs that are involved in pre-mRNA splicing, were over-represented in up-regulated sRNAs on drought. Alternative splicing of pre-mRNA splicing under drought stress was observed in multiple tissues, particularly in the leaf and ear (Thatcher *et al*. 2016), which might partially attributes to amount and stability of various splicing sn-RNAs.

The snoRNAs primarily include two classes of sRNAs, box C/D and box H/ACA snoRNAs, which guide methylation and pseudouridylation of other RNAs, respectively (Bachellerie *et al*. 2002; Kiss 2006). The snoRNA-mediated chemical modifications of rRNAs and splicing snRNAs have been demonstrated to be essential for ribosomal function as well as mRNA splicing and maturation (Morris and Mattick 2014; Dupuis-Sandoval *et al*. 2015). The sno-sRNA was over-represented in both down-regulated and transiently down-regulated sRNA groups under drought stress. Down-regulation of sno-sRNAs may be the result of the reduction of snoRNAs, which would reduce the activity of methylation and pseudouridylation of rRNAs and splicing snRNAs. Given the reduction of sno-sRNAs and the increase of rsRNA and splicing sn-sRNAs upon drought stress, it is tempting to speculate that rRNAs and splicing snRNAs are destabilized with decreased methylations or pseudouridylations as mediated by snoRNAs. Both changes in chemical modification, presumably, and the quantity of rRNAs upon drought stress could effectively alter the activity of the protein synthesis machinery. The observation of sRNA changes related to rRNAs and splicing snRNAs indicates the post-transcriptional regulation is an important mechanism for adaptive response to drought stress. snoRNAs exhibiting responses to drought were also found in another plant species (Hackenberg *et al*. 2015). Recently, snoRNAs were also found to be involved in metabolic stress responses, including oxidative stress in human cells (Michel *et al*. 2011; Chu *et al*. 2012; Youssef *et al*. 2015). Taken together, we propose that snoRNAs play a role to regulate biological processes under drought stress through chemical modifications of rRNAs and splicing snRNAs.

## Acknowledgements

This work was supported by the National Key Research and Development Program of China (Grant No. 2016YFD0101002), the National Natural Science Foundation of China (Grant No. 31330056), the Agricultural Science and Technology Innovation Program of CAAS, and the U.S. National Science Foundation (Award No. 1238189). This work of Haiyan Wang was partially supported by a grant from the Simon Foundation (Award# 246077). We thank the support from the Kansas Agricultural Experiment Station (contribution No. 17-132-J).

## Authors’ contributions

JZ and GW designed the study. JZ, YD, ZJ and KW performed experiments and generated the sequence data. SL, EZ, CH, JF, YH, ML, WL, and HW analyzed data. SL, FW, JZ, and EZ wrote the manuscript. FW, HW, EZ, WL, and GW revised the manuscript. All authors reviewed and approved the final manuscript.

